# The C-terminal domain of SEC-10 is fundamental for exocyst function, Spitzenkörper organization and cell morphogenesis in *Neurospora crassa*

**DOI:** 10.1101/2022.01.14.475644

**Authors:** Alfredo Figueroa-Meléndez, Leonora Martínez-Núñez, Adriana M. Rico-Ramírez, Juan M. Martínez-Andrade, Mary Munson, Meritxell Riquelme

## Abstract

The exocyst is a conserved multimeric complex that participates in the final steps of the secretion of vesicles. In the filamentous fungus *Neurospora crassa,* the exocyst is crucial for polar growth, morphology, and the organization of the Spitzenkörper (Spk), the apical body where secretory vesicles accumulate before being delivered to the plasma membrane. In the highly polarized cells of *N. crassa,* the exocyst subunits SEC-3, SEC-5, SEC-6, SEC-8, and SEC-15 were previously found localized at the plasma membrane of the cells’ apices, while EXO-70 and EXO-84 occupied the frontal outer layer of the Spk, occupied by vesicles. The localization of SEC-10 had remained so far elusive. In this work, SEC-10 was tagged with the green fluorescent protein (GFP) either at its N- or C-terminus and found localized at the plasma membrane of growing hyphal tips, similar to what was previously observed for some exocyst subunits. While expression of an N-terminally tagged version of SEC-10 at its native locus was fully viable, expression of a C-terminally tagged version at its native locus resulted in severe hyphal growth and polarity defects. Additionally, a *sec-10* knockout mutant in a heterokaryotic state (with genetically different nuclei) was viable but showed a strongly aberrant phenotype, confirming that this subunit is essential to maintain hyphal morphogenesis. Transmission electron microscopy analysis revealed the lack of a Spk in the SEC-10-GFP strain, suggesting a critical role of the exocyst in the vesicular organization at the Spk. Mass spectrometry analysis revealed fewer peptides of exocyst subunits interacting with SEC-10-GFP than with GFP-SEC-10, suggesting an essential role of the C-terminus of SEC-10 in exocyst assembly and/or stability. Altogether, our data suggest that an unobstructed C-terminus of SEC-10 is indispensable for the exocyst complex function and that a GFP tag could be blocking important subunit-subunit interactions.

## Introduction

Filamentous fungi are excellent model organisms to study polarized growth. Fungal hyphae extend by a sustained polarized process that involves the apical exocytosis of secretory vesicles that fuse with the plasma membrane (PM), allowing the secretory vesicles to deliver their content to the extracellular space (Riquelme et al., 2011; Rizzoli and Jahn, 2007). Exocytosis is fundamental for many cellular activities, including neurotransmitter release at the presynaptic terminal, vesicular transport to the basolateral membrane, and primary ciliogenesis in animal cells, expansion of root hair tips in plants, and hyphal tip growth in fungi, all of which involve polarized secretion of vesicles (Grindstaff et al., 1998; Kennedy and Ehlers, 2011; Lopez-Franco et al., 1994; Monshausen et al., 2008; Zuo et al., 2009). The interaction between a vesicle and a target membrane throughout the different steps of the secretory and endocytic pathways is mediated by multisubunit tethering complexes (MTCs) (Dubuke and Munson, 2016). The exocyst complex is a highly conserved MTC within eukaryotes that was first identified in the budding yeast *Saccharomyces cerevisiae.* It consists of the proteins Sec3, Sec5, Sec6, Sec8, Sec10, Secl5, Exo70, and Exo84 (Guo et al., 1999; TerBush et al., 1996; TerBush and Novick, 1995). This complex has been linked to several diseases and exocyst mutants are associated with cell growth and developmental defects, as has been shown in mouse and *Drosophila* models (Friedrich et al., 1997; Martin-Urdiroz et al., 2016; Murthy et al., 2003; Murthy et al., 2005). Consistent with an essential role in fusion of secretory vesicles at exocytic sites, temperature sensitive mutants of exocyst subunits cause an accumulation of secretory vesicles in the cytoplasm (Novick, 1980).

The exocyst is proposed to mediate the tethering of post Golgi secretory vesicles to the PM and to promote the fusion of membranes via SNARE complexes (Heider and Munson, 2012; TerBush et al., 1996; Wu and Guo, 2015; Yue et al., 2017), but mechanistic details are lacking. A reconstruction of the 3D architecture of the S. *cerevisiae* exocyst *in vivo* suggested that the subunits’ N-terminal ends are oriented towards the center of the complex, except for Sec10, whose C-terminus is the one oriented toward the center (Picco et al., 2017). The near-atomic cryo-EM structure of the S. *cerevisiae* exocyst revealed that all exocyst subunits are rod-shaped (Lepore et al., 2018; Mei et al., 2018). The subunits contain long N-terminal (except for Sec3, where the region is centrally located) coiled regions called CorEx (core of the exocyst) that assemble together to generate the framework of the complex in the form of two modules of four subunits each, which consist of subcomplex I (Sec3, Sec5, Sec6 and Sec8) and subcomplex II (Sec10, Sec15, Exo70 and Exo84) that pack against one another (Lepore et al., 2018; Mei et al., 2018).

In the filamentous fungus *Neurosporo crassa,* a full exocyst with the eight subunits is required for maintaining polarized hyphal growth and for the orderly arrangement of large secretory vesicles (macrovesicles) at the Spitzenkörper (Spk) (Riquelme et al., 2014). Deletion of *sec-5* severely affected cell morphogenesis and the apical localization of macrovesicles in *N. crassa.* GFP tags were added at the 3’ end of the endogenous loci encoding the different exocyst subunits; the subunits SEC-5, SEC-6, SEC-8, and SEC-15 localized at the PM in *N. crassa* hyphal tips, whereas EXOmicrovesicles, and another strain that expresses-70 and EXO-84 localized at the frontal periphery of the Spk (Riquelme et al., 2014). SEC-3 was observed at both the PM and the Spk. In the *sec-5* mutant background, SEC-6-GFP and EXO-70-GFP were not localized at the hyphal apex, suggesting that SEC-6 and EXO-70 localization depends on SEC-5 (Riquelme et al., 2014).

Curiously, in previous studies, no viable transformed *N. crassa* strains could be recovered for the C-terminally GFP tagged SEC-10 subunit. Similar results were reported for the rice blast fungus *Magnaporthe oryzae,* where SEC-10 was the only subunit that could not be successfully tagged, which suggested that a tag at the C-terminus of SEC-10 could interfere with function in SEC-10 nearby subunits, or exocyst binding partners (Gupta et al., 2015). Further, S. *cerevisiae* cells overexpressing SeclOΔC, a protein lacking the C-terminal region, led to a block in exocytosis and accumulation of vesicles (Roth et al., 1998).

In this study, we endogenously tagged the SEC-10 exocyst subunit with GFP at either its N-or C-terminus, as well as produced mutants of *N. crassa* either lacking *sec-10* or harboring a truncated version of *sec-10,* to test the importance of the C-terminal region. We analyzed by SDS-PAGE and mass spectrometry the proteins recovered from pull downs of both the C-terminally tagged SEC-10 and the N-terminally tagged SEC-10. We found that deleting *sec-10* or tagging *sec-10* at the C-terminus had a similar detrimental effect on growth, whereas tagging *sec-10* at the N-terminus or eliminating 40 amino acids from the C-terminus of sec-10 had no significant impact on growth rate. Our results suggest that the GFP tag at the C-terminus affects the stability of the complex by acting as a spatial impediment for the assembly of the exocyst subunits and could be destabilizing important protein-protein interactions within the complex.

## Materials and methods

### Strains used or generated in this study

The *N. crassa* strain FGSC9718 was used to tag SEC-10 with GFP and to delete the *sec-10* gene. All strains were cultured in Vogel’s Minimal Medium (VMM; (Vogel, 1956) supplemented with 1.5% sucrose and incubated at 30 °C. To select putative transformants, VMM was supplemented with 0.05% fructose, 0.05% glucose, 2% sorbose (FGS), and hygromycin B (Hyg, 300 μg/mL; Invitrogen, Carlsbad, CA). All *N. crassa* strains that were used or developed in this study are listed in Table 1.

### *In silico* analysis of SEC-10

Functional and structural information about SEC-10 was analyzed using the UniProt protein identification code for *N. crassa* (Q7SD81_NEURCR) on several different servers. Similar analyses were performed for SEC-10 orthologues found in *C. albicans, S. cerevisiae, A. nidulans* and *M. oryzae.* The different functional domains were identified using the UniProtKB database (https://www.uniprot.org/). The motifs contained in these sequences were identified using ScanProsite (https://prosite.expasy.org) and superfamily domains on the InterPro server (http://www.ebi.ac.uk/interpro/). The presence of predicted disordered domains in the subunits’ structure was detected with the MobiDB server, which uses a consensus method that is optimized to find long intrinsically disordered protein (IDRs). MobiDB is integrated in InterProScan and its predictions are propagated to several EBI resources (PDBe, UniProt, InterPro). (http://mobidb.bio.unipd.it/).

### Tagging SEC-10 with GFP

The SEC-10 protein was endogenously tagged at the C- or N-terminus using the Split-Marker gene replacement technique as previously described (Smith et al., 2011). The C-terminus was tagged by using constructs for homologous gene replacement based on pGFP::hph::loxP (Honda and Selker, 2009); GeneBank accession number FJ457011; Fig. S1A), whereas the N-terminus was tagged by using constructs based on pGFP::hph::loxP and pCCG::N-GFP (Honda and Selker, 2009); Fig. S1B). The replacement cassettes contained the selectable hygromycin resistance marker *(hph)* (Fig. S1B). The genomic regions of *sec-10* were amplified from genomic DNA of *N. crassa* N1 that was extracted with a DNeasy Plant Mini Kit (Qiagen, Valencia, CA) (Table 1). A total of 1 μg of DNA (500 ng of each DNA replacement cassette) was mixed with 1.25E8 macroconidia of the FGSC9718 strain and transferred to 0.2 cm-gap sterile electroporation cuvettes (BIO-RAD^®^ GenePulser Xcell™) for transformation by electroporation (1500 V, 25 μF, 600 Ohm) (Margolin et al., 1997)). Electroporated cells were plated in VMM-FGS agar containing a final concentration of 300 μg/mL of hygromycin. SEC-10 was also tagged ectopically at the C-terminus by using a *his-3* targeting vector containing a *Pccg1::sec-10::gfp* cassette (VMRP-114; unpublished). The plasmid (1 μg) linearized with *Nde*l and treated with Shrimp Alkaline Phosphatase was used to transform 1.25E8 macroconidia of the FGSC9717 histidine auxotrophic strain via electroporation. The genetic constructs and the fusions are shown in Supplementary figure 1.

### Laser scanning confocal microscopy cell preparations

Conidia from a preserved stock were inoculated on VMM 1.5% agar plates and incubated overnight at 30 °C. A small rectangular section of 1.5 cm x 2 cm was cut out from the plate with a sterile spatula. The face of the agar containing the mycelium was placed carefully over a cover slide (VWR VistaVision™ Cover Glasses 16004-096, No. 1, 24 x 60 mm) and left at room temperature for 15 min. A small drop of immersion oil was placed over a 60X objective lens. The slide was loaded on the stage on an Olympus FluoView™ FV1000 laser scanning confocal microscope (LSCM). The images were acquired on a confocal LSM observation mode with a PLAPON 60X O NA:1.42 objective lens, a oneway XYT scan mode, and a scanning speed of 20 us/pixel. The region of interest was clipped with a rectangular tool of the FLUOVIEW FV1000 software version 4.0.2.9. To image GFP, a blue Argon-ion laser was used with an excitation wavelength of 488 nm and an emission wavelength of 510 nm. In the case of cells expressing the mCherry fluorescent protein (mChFP) or stained with FM4-64, a green Helium-Neon laser was used with an excitation wavelength of 543 nm and an emission wavelength of 612 nm. FM4-64 is a lipophilic stain that labels membranes inside the cell (Pogliano et al., 1999). Images of the colonies were taken to compare the phenotype and characteristics of the strains. The strains were incubated 24 h at 30 °C, and photographs of these colonies were taken (Nikon Digital Camera D33697, Nikon Corp., Japan). Each colony was observed under a stereomicroscope (Olympus Optical Co., Ltd. Model SZX-ILLB2-100) on different objective lenses.

### Transmission electron microscopy cell preparations

Conidia of *N. crassa* strain expressing SEC-10-GFP were inoculated on sterile and deionized dialysis membranes overlaid on VMM 1.5% agar plates and grown at 27 °C for 48 h. Subsequently, 5×5 cm squares (n=80) containing growing hyphae were cut, let recover for 30 min, and cryo-fixed by rapidly immersing them in liquid propane cooled to −186 °C with liquid nitrogen as previously described (Dunn and Wobbe, 1993). The samples were transferred to 2% osmium tetroxide and 0.05% uranyl acetate in acetone at −80°C for 72 h. After that, they were washed three times in 100% anhydrous acetone, infiltrated in Spurr resin, and embedded in Teflon coated glass slides to be polymerized at 60 °C for 24 h. For semi-quantitative analysis, hyphae (n=10) were selected randomly from different regions of the samples to obtain ultrathin medial sections (70 nm). All the samples were observed using a Hitachi H7500 transmission electron microscope to 80 kV with a 16-megapixel Gatan CCD digital camera.

### Producing *sec-10* mutant strains

An *N. crassa sec-10* knock-out mutant strain was obtained from the Fungal Genetic Stock Center (FGSC11723) and tested to confirm the gene’s deletion. Genomic DNA was extracted and tested via PCR with oligonucleotides that flank the *sec-10* open reading frame (ORF). Since an amplicon of the corresponding *sec-10* ORF length was obtained from the presumable mutant, confirming a heterokaryotic stage, recovery of a homokaryotic strain was attempted by genetic crossing with the wild-type strain N150 (FGSC9013). Heterokaryosis refers to the presence of genetically distinct nuclei within the same cell, and homokaryosis refers to the presence of genetically identical nuclei within the cell (Strom and Bushley, 2016). However, deletion of the *sec-10* ORF could not be confirmed through PCR. These results led us to produce our own *sec-10* knock-out mutant strain by using the Split-Marker gene replacement technique (Fig. S1C). Genomic DNA from *N. crassa* Wild Type FGSC988 was extracted with a DNeasy Plant Mini Kit (Qiagen, Valencia, CA). The plasmid pGFP::hph::loxP was purified with a Qiagen plasmid isolation kit. The knock-out constructs were produced using genomic DNA and oligonucleotides with *hph* gene sequence overhangs (Table 2). The 1 kb of the 5□ untranslated region (UTR) was amplified with forward primer 720 and reverse primer 721 that flank the *hph* gene. The 1 kb of the 3□ UTR was amplified with a forward primer that contains an overhang that overlaps with the 5’ region of the *hph* gene and a reverse primer that overlaps with the 3’ end of 3’ UTR region. These two constructs flank the ORF region of *sec-10,* and upon homologous recombination the *hph* gene replaces *sec-10* (Fig. S1C). A deletion mutant named *sec-10^840_880del^* was also created to test the effects of deleting the C-terminal disordered domain, which consists of 40 amino acids by eliminating the last 120 nucleotides of the *sec-10* ORF (Fig. S1D), which corresponds to a short C-terminus disordered domain. The reverse oligonucleotide included an overhang that overlaps with the *hph* gene and excluded the 120 nucleotides by moving the stop codon 120 nucleotides upstream (Fig. S1D). *N. crassa* strain FGSC9718 macroconidia were transformed with the DNA constructs by electroporation. Transformants were plated on VMM agar solidified with hygromycin at a final concentration of 300 μg/mL.

### SEC-10-GFP and GFP-SEC-10 pull-downs with Lag 94-15 for mass spectrometry analysis

Strains expressing cytoplasmic GFP, N-terminal tagged SEC-10, and C-terminal tagged SEC-10 were grown on VMM plates. The mycelium was scraped from the agar and transferred to Erlenmeyer flasks with liquid VMM for incubation at 200 rpm, 30 °C (INFORS Multitron CH-4103). After vacuum filtration, the recovered biomass was pulverized in a planetary ball mill (Retsch PM100) under liquid nitrogen conditions and stored at −80 °C until use. 150 mg of frozen powder were added to a 1.5 mL microcentrifuge tube and resuspended in 650 μL lysis buffer (20 mM Tris pH 8.5, 150 mM KCl, 0.1% Tween 20 and 1x cOmplete Mini EDTA-free protease inhibitor solution; Roche Life Science). The suspension was spun at 14,000 g for 10 min at 4°C. The supernatant was incubated with 15 μL of M270 magnetic amine Dynabeads covalently bound to anti-GFP LaG94-15 nanobody (Fridy et al., 2014) at 4 °C for 1 h, nutating. The beads were washed three times with lysis buffer and resuspended in 25 μL of lysis buffer and 5 μL of 5X SDS-PAGE buffer. The samples were boiled at 95 °C for 5 min and used to run an SDS-PAGE (Novex™ WedgeWell™ 4 to 20%, Tris-Glycine, 1.0 mm, Mini Protein Gel) followed by 1X Krypton™ fluorescent staining (Thermo Fischer Scientific). Proteins were visualized on a Typhoon FLA 9000 (Ex/Em = 532/580 nm).

### Mass spectrometry sample preparation

For mass spectrometry analysis, the pull-down was scaled up to 750 mg of pulverized mycelium resuspended in 1 mL of lysis buffer. The sample was run a small distance into the gel by electrophoresis so that the gel sample would contain the exocyst complex binding partners. The gel was stained with Coomassie solution and cut into 1×1 mm pieces per sample, which were kept in 1.5 mL tubes with milliQ. water until mass spectrometry analysis. The water was discarded, and the gel pieces were incubated in 10 mM 1,4 dithiothreitol (DTT) in ammonium bicarbonate solution at 50 °C to reduce the disulfide bands, followed by iodoacetamide (IAA) alkylation at room temperature. The remaining DTT and IAA were removed, followed by three wash steps with water, 50 mM ammonium bicarbonate: acetonitrile (50:50), and 100% acetonitrile. The acetonitrile was discarded, and the samples were dried in a Speed Vac lyophilizer. After complete drying, trypsin enzyme and proteaseMAX surfactant in ammonium bicarbonate solution were placed in the tubes, and digestion was performed for 18 hours at 37 °C. Then the samples were spun down, the supernatant was removed and placed in a new tube, and gel pieces were further dehydrated in a solution of acetonitrile: 1% formic acid in water (80:20) while. The samples were spun down, and the resulted supernatant was combined with the previous one, and the peptide mixture was dried down in a Speed Vac lyophilizer. The obtained peptide pellets were resolubilized in 5% acetonitrile with 0.1% formic acid in water to prepare the samples for analysis.

### Liquid chromatography and mass chromatography analysis

The liquid chromatography tandem mass spectrometry (LC-MS/MS) analysis was performed on a Waters nanoAcquity UPLC (Waters, Milford, MA) and a Q. Exactive hybrid quadrupole-Orbitrap mass spectrometer (Thermo Fisher Scientific Inc., Waltham, MA).

The peptides were trapped (4 mins) in an in-house packed pre-column (C18, 200A, 5μm, 2cm), then the peptide separation started at 5% organic phase B (acetonitrile with 0.1 % formic acid) and 95% aqueous phase A (water with 0.1 % formic acid). The gradient was carried on to 35% B over 90 mins, on an inhouse packed analytical column (C18, 100A, 3 μm, 25 cm), followed by 90% B for 10 mins wash, and 5% B for 15 mins (re-equilibration). The MS data acquisition was performed in positive electrospray ionization mode (ESI+), within the mass range of 300-1750 Da for MS1, with the orbitrap resolution of 70,000 at m/z 200. The maximum injection time and the AGC target were set to 30 ms and 1e6, respectively. Data Dependent Acquisition (DDA) MS/MS was applied on the top 10 precursor ions within a 1.6 Da isolation window, with the resolution of 17500 at m/z 200. The maximum injection time and the AGC target for the MS/MS were set to 110 ms and 1e5, respectively. The normalized collision energy was set to 27 V for fragmentation.

### Mass spectrometry data processing

The acquired MS raw data was imported to Thermo Proteome Discoverer (PD) 2.1.1.21 (Thermo Fisher Scientific Inc.) software for processing. The data was searched against UniProt database using Mascot Server 2.6.2 (Matrix Science Ltd) in PD. Two maximum missed cleavages were considered for trypsin digestion. The peptide modifications were set to carbamidomethyl of cysteine as fixed modification, and three variable modifications of methionine oxidation, N-terminal acetylation, and N-terminal glutamine to pyroglutamate. The error tolerance was specified as 10 ppm for the monoisotopic mass of the precursor and 0.05 Da for the fragment mass. The data was further processed by Scaffold 4.10.0 (Proteome Software Inc.) to validate peptide and protein identification. Scaffold applied a 1% false discovery rate (FDR) for peptide identification based on the peptide Prophet algorithm (Keller et al., 2002). A minimum number of 2 peptides was considered as the threshold for protein identification with 99% probability, applied by Scaffold, using protein Prophet algorithm (Nesvizhskii et al., 2003).

## Results

### The exocyst subunit SEC-10 localizes at the plasma membrane in hyphal tips

*N. crassa* SEC-10 orthologues in *Saccharomyces cerevisiae, Magnaporthe oryzae, Aspergillus nidulans,* and *Candida albicans* share a similar length, ranging from 785 to 880 amino acids. All SEC-10 orthologues contain a predicted coiled-coil domain or domains, which are ~100 amino acids downstream from the N terminus (Fig. 1). Previously, SEC-10 was the only exocyst subunit that could not be successfully tagged endogenously at its C-terminus in *N. crassa* (Riquelme et al., 2014). In this study, we successfully obtained an *N. crassa* strain that expressed SEC-10 tagged with GFP. We created strains in which SEC-10 was both C-terminally (SEC-10-GFP) and N-terminally (GFP-SEC-10) tagged with GFP at its native locus, under the control of the native promoter (Fig. 2A), and also C-terminally tagged at the *his-3* locus under the control of the *ccg-1* promoter (pCCG-1-SEC-10-GFP; Fig. 2B). In all three cases, the resulting GFP-tagged SEC-10 protein localized at the apical PM (Fig. 2), similar to the localization previously described for the exocyst subunits SEC-5, SEC-6, SEC-8, and SEC-15 (Riquelme et al., 2014). Localization of GFP-SEC-10 fluorescence was similar to localization of SEC-10-GFP. Neither SEC-10-GFP nor GFP-SEC-10 showed co-localization with the Spk at the tip, as stained in live cells with FM4-64 (Fig. 2C, E). In *N. crassa,* we can produce both homokaryon strains, which contain genetically identical nuclei, and heterokaryons, which contains two genetically different nuclei. We found that the homokaryon strain expressing SEC-10-GFP had irregular cell shapes, a barely visible Spk and a clear problem with maintaining growth (Fig. 2D).

**Figure 1.**
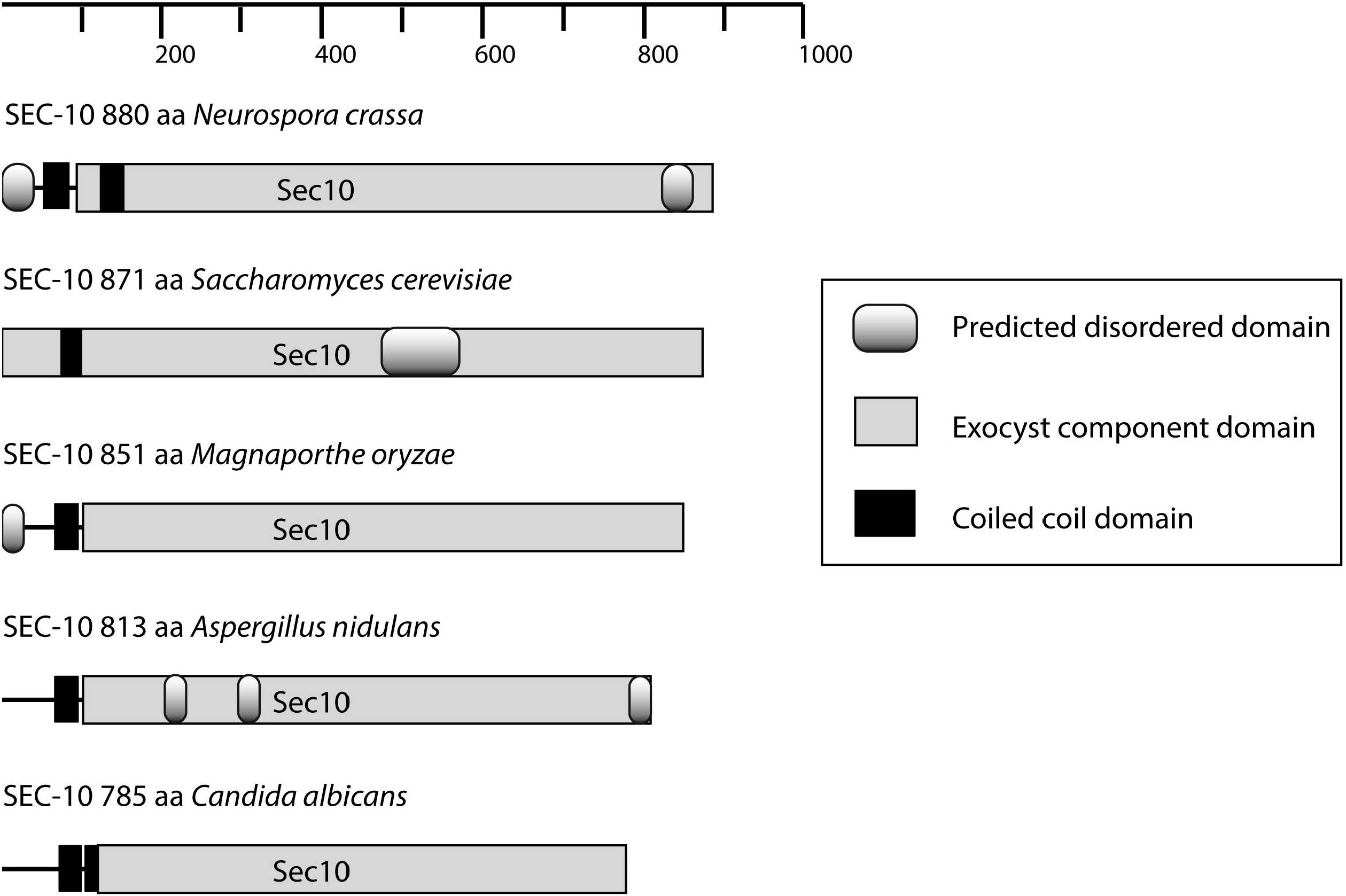
SEC-10 orthologues in other fungi have similar sizes and a conserved exocyst component sequence. Scale bar indicates amino acid residues in the sequence of the orthologues and ranges from 785 aa to 880 aa. Predicted domains found in SEC-10 orthologues are shown in the box legend; the size of the domain represents the actual scale of the sequence.

**Figure 2.**
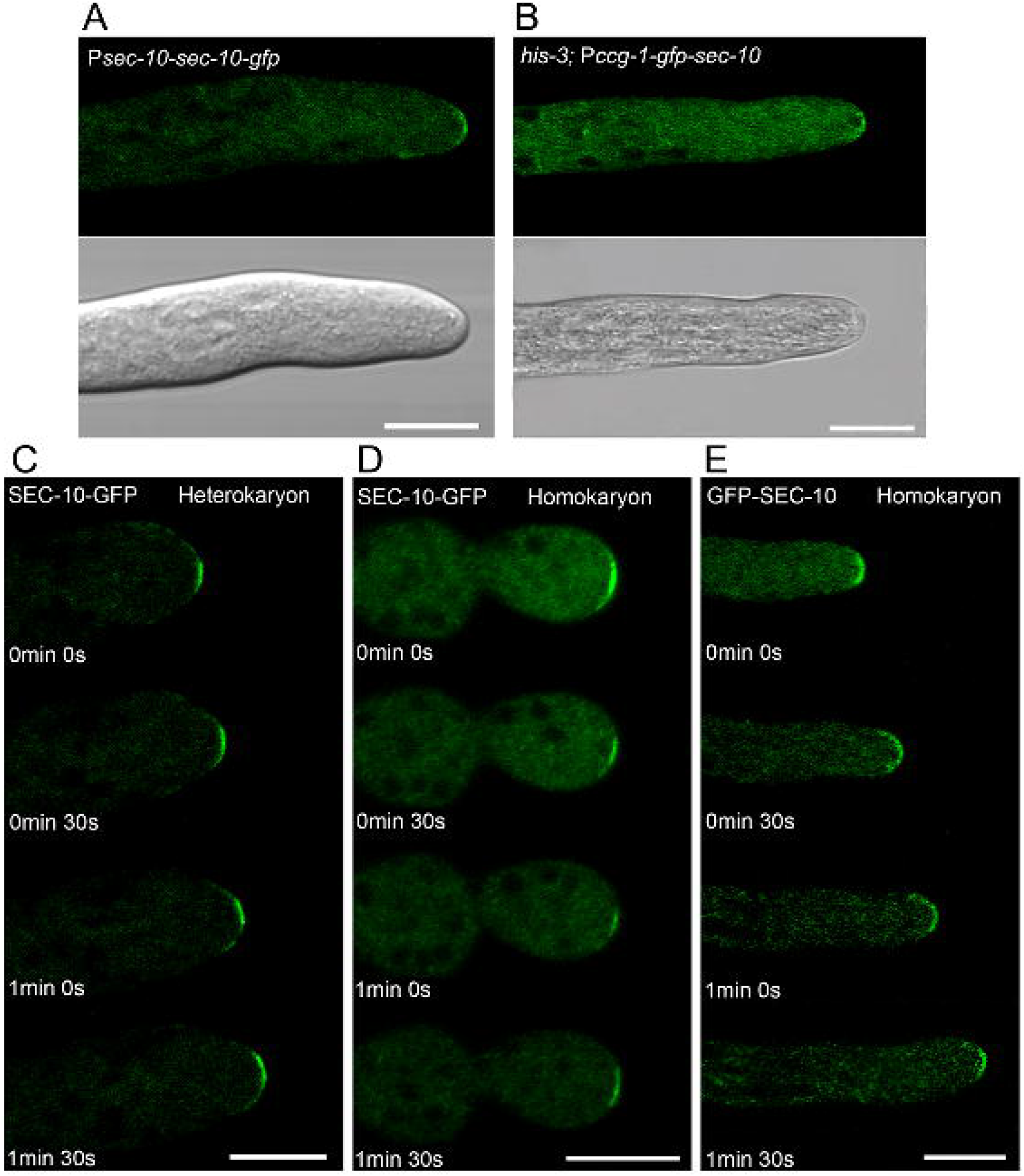
Laser scanning confocal microscopy of *N. crassa* hyphae expressing SEC-10-GFP and GFP-SEC-10 reveal that SEC-10 protein is localized at the hyphal tip in the apical dome: A) Under control of native promoter and in a heterokaryotic state, and B) Under control of *Pccg-1* expressed ectopically from the *his-3* locus. C) Time lapse of growing hypha that expresses SEC-10-GFP in a heterokaryotic state. D) Time lapse of SEC-10-GFP expressed in a homokaryotic state. E) Time lapse N-terminal GFP-SEC-10 expressed in a homokaryotic state. Scale bars = 10 μm.

### GFP tagging of SEC-10 at the C-terminus caused growth defects and hyphal deformities

We compared the growth rates and cellular phenotypes of the endogenously tagged SEC-10-GFP heterokaryon, SEC-10-GFP homokaryon, GFP-SEC-10 homokaryon, and wild type 9718 strains. The corresponding genetic constructs introduced in each strain are shown in Fig. 6A. Growth defects and cell deformities were detected in the SEC-10-GFP homokaryon strain (Supplementary video 1) and to a lesser degree in the SEC-10-GFP heterokaryon strain. Differential interference contrast (DIC) microscopy and Laser scanning confocal microscopy (LSCM) revealed that the homokaryon strain SEC-10-GFP had a dispersed accumulation of FM4-64 at swollen hyphal tips (Fig. 3A), and a larger number of septa, resulting in unusually shorter compartments (Fig. 3B; 26 ± 0.8 μm, N=15) when compared to the wildtype strain (Fig 3C; 159 ± 10.2 μm, N=15). After 24 h of growth, the wild type, and the C-terminal SEC-10 tagged strains showed clear differences in colony growth (Fig. 4). The wild-type strain covered almost the entire agar surface of the Petri dish, whereas the SEC-10-GFP heterokaryon had only begun to colonize the surface, and in the homokaryotic state, the SEC-10-GFP strain had not yet developed. It was only after 7 days of growth when the SEC-10-GFP heterokaryon had fully colonized the surface. The homokaryon colony did not grow any further along the medium but began growing aerially (Fig. 4A). Stereomicrographs of these strains highlighted how compact the mycelium of the SEC-10-GFP homokaryon strain was when compared to the other strains (Fig. 4B). DIC microscopy revealed irregular cell shapes in all observed hyphae in both the SEC-10-GFP homokaryon and heterokaryon strains, although more so in the homokaryon (Fig. 4C; Fig. 6B). Notably, the hyphae of the SEC-10-GFP homokaryon strain displayed a larger hyphal diameter (11.8±0.7 μm, N□=□25) than the wild-type strain (Fig. 4C; 8.46±0.7 μm, N□=□25) and the GFP-SEC-10 homokaryon strain (7.6±0.6 μm, N□=□25). Another characteristic of this strain was how the hyphae change direction momentarily, with a concomitant loss of the Spk localization (Fig. 5.; Supplementary video 2).

**Figure 3.**
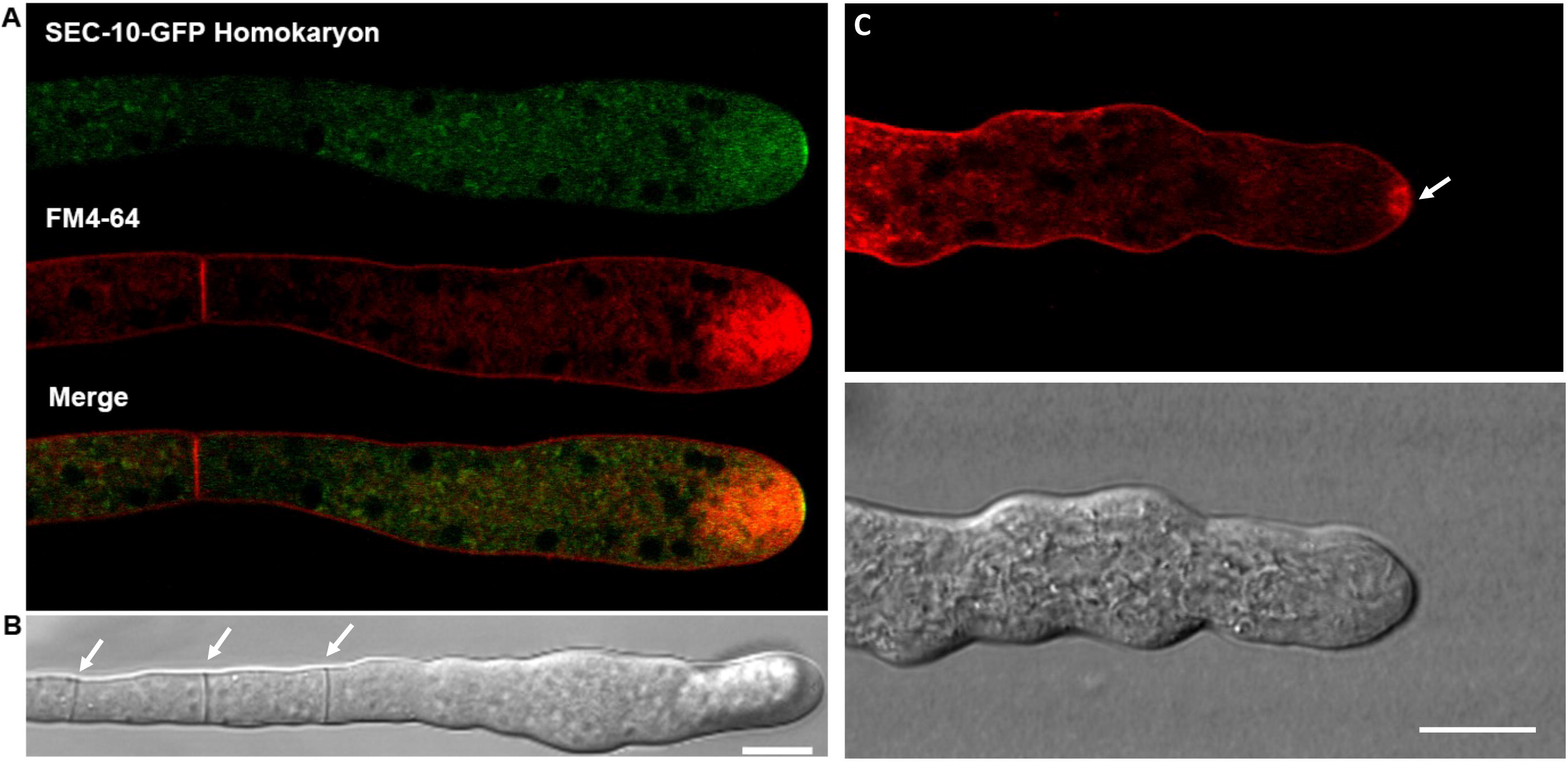
DIC and LSCM of SEC-10-GFP homokaryon strains show hyphae that are hyperseptated and swollen at the tips. A) FM4-64 staining of the SEC-10-GFP strain shows that FM4-64 stained the plasma membrane surrounding the cytoplasm, the septa and the accumulated at the apex in a disperse cloud. B) Differential interference contrast (DIC) microscopy of a hypha of the SEC-10-GFP homokaryon strain. Under DIC the hypha revealed an unusual pattern of subapical septa (white arrows). In contrast, C) FM4-64 staining of a wild-type strain showing the stained plasma membrane and the Spk (white arrow) with an unstained core (top image). DIC microscopy of the same hypha displayed above (bottom image). Scale bars = 10 μm.

**Figure 4.**
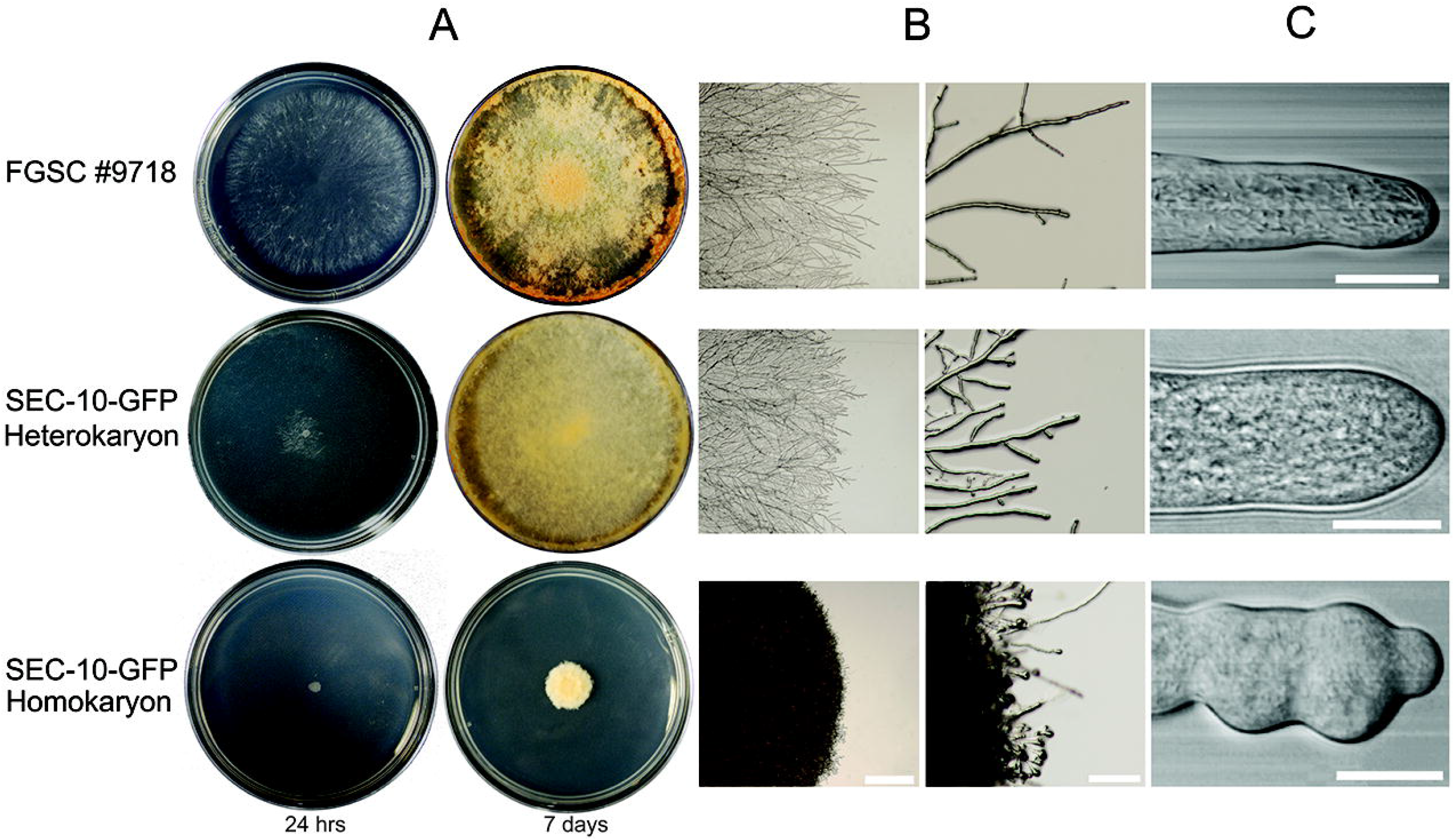
Qualitative comparison of the growth and morphology of the strains expressing SEC-10-GFP heterokaryon and homokaryon and the wild-type strain 9718. A) Phenotype of the different strains of *N. crassa* grown on MMV plates and incubated for 24 hours and for one week at 30 ° C. B) Morphology of the edge of the colonies viewed by stereomicroscopy. C) Differential interference contrast (DIC) microscopy of hyphae observed with a 6OX objective. Scale bars B = 2mm and 200 μm, C = 10 μm

**Figure 5.**
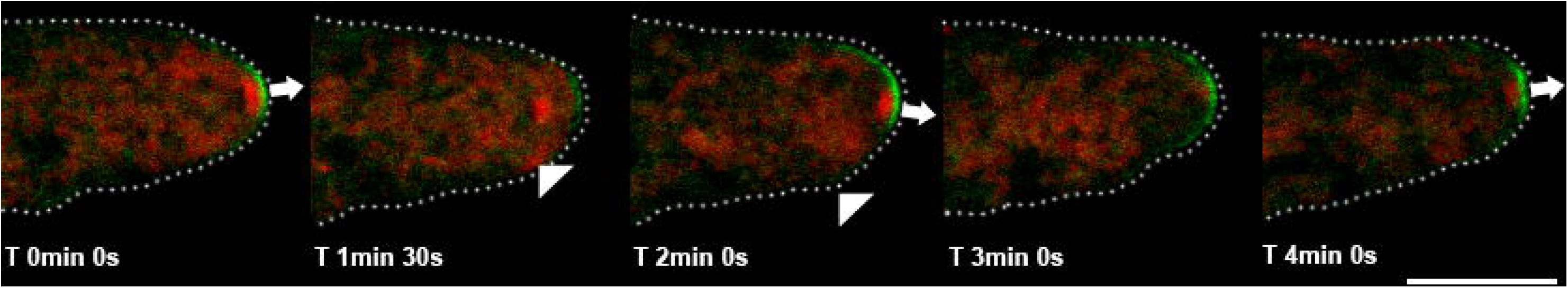
Effect of the expression of SEC-10-GFP in a heterokaryon strain on Spk organization and growth revealed by time-lapse microscopy. Spk was stained with FM4-64 and observed by Laser scanning confocal microscopy. SEC-10-GFP heterokaryons have minor irregularities in cell shape, as seen in the white dotted outline of the cells. The fluorescence of the Spk is lost as growth pauses and reappears when growth resumes. The position of the FM4-64 stained Spk changes with the corresponding changes in growth direction (white arrows). Scale bar = 10 μm.

**Figure 6.**
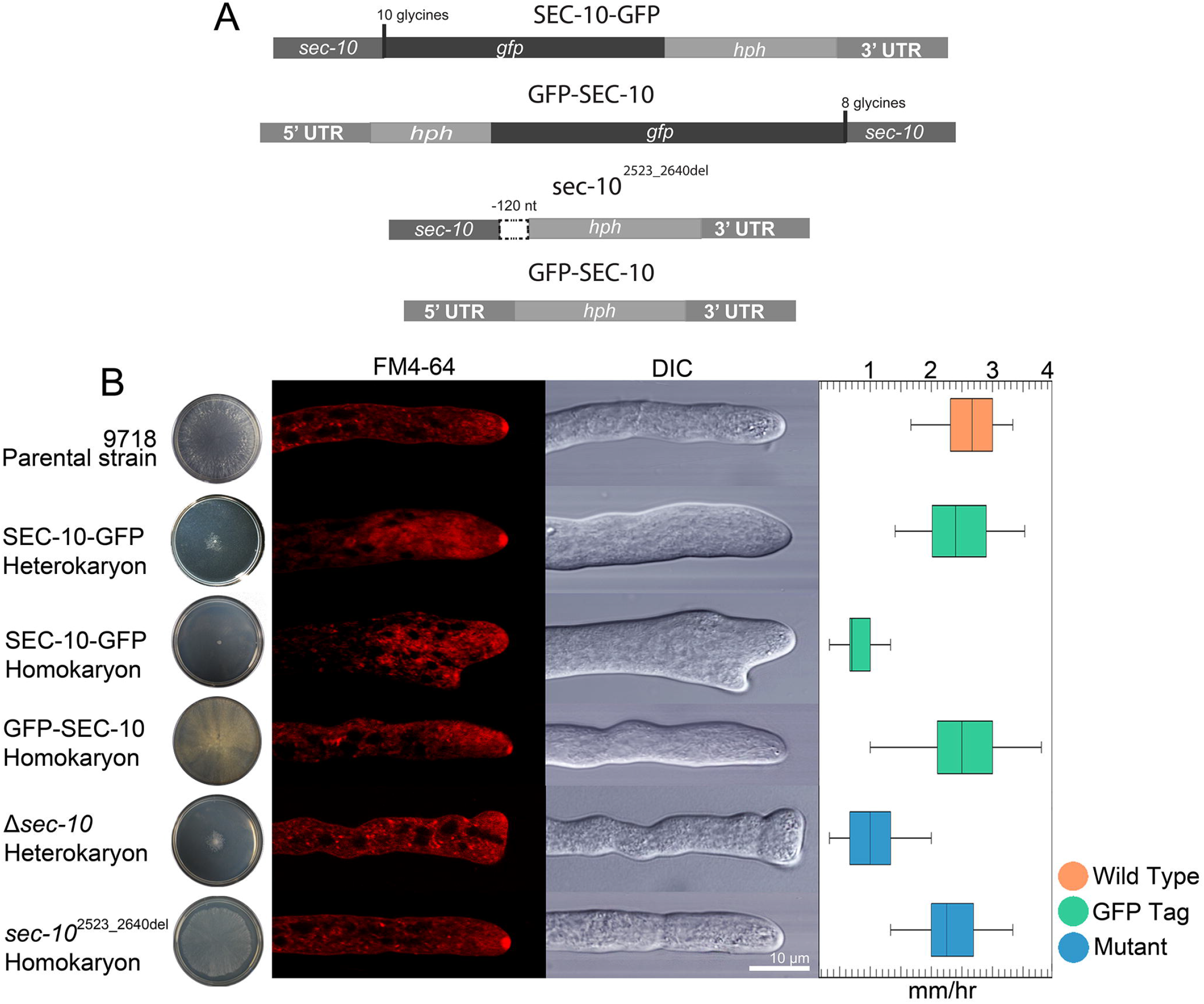
Tagging SEC-10 with GFP at its C-terminus disrupts growth in homokaryons, and the *sec-10* knock out mutation is lethal and dominant in heterokaryons. A) Schematic view of the DNA constructs used. DNA constructs were incorporated into endogenous loci via electroporation and the strains were selected by hygromycin resistance *(hph).* B) Left column shows colony growth of the strains after 24 hours. Laser scanning confocal microscopy of strains stained with FM4-64 reveal vesicle distribution inside the cell. The DIC microscopy shows the difference in morphology between the different strains. The right column chart shows growth rate of the strains: wild type (orange), GFP tagged (green), and *sec-10* mutants (blue). Scale bar = 10 μm.

### *sec-10* is an essential gene required for cell growth and morphogenesis

We next examined the phenotype of strains in which *sec-10* was deleted or truncated. The *Δsec-10* mutant could not be obtained in a homokaryotic state by either microconidia production or genetic crossing, indicating that it is essential. However, the heterokaryon was viable but with delayed growth and almost a complete lack of conidia production; this mutant strain had a 99% reduction in macroconidia production, and 95% of the conidia produced were microconidia. In contrast, strain *sec-* 10^840_880del^, θ mutant with a c-terminally truncated SEC-10 protein, had no phenotypic or growth defects (Fig. 6B). Upon comparing the growth rate of the strains 9718 (2.8±0.29μm, N□=□16), SEC-10-GFP heterokaryon (0.76±0. 0.39μm, N□=□16) and homokaryon (0.16±0.01μm, N□=□16), GFP-SEC-10 homokaryon (2.4±0.45μm, N□=□16), *Δsec-10* heterokaryon (0.9±0.29μm, N□=□16), and sec-10^840_880del^ homokaryon (2.5±0.37μm, N□=□16) (Fig. 6B), it was revealed that the growth rates of the C-terminal tagged SEC-10 homokaryon strain and the *Δsec-10* heterokaryon mutant strain were significantly reduced. In contrast, the N-terminal tagged SEC-10 strain did not show a significantly different growth rate than the wild-type strain. Together, our finding that the SEC-10-GFP homokaryon strain had an 83% growth rate reduction, as well as morphologic and cell polarity defects, indicates that the GFP tag at the C-terminus of SEC-10 perturbs exocyst function.

The genetic constructs that were used for electroporation are shown in Figure 6A, including the *sec-10* mutants Δ*sec-10* and *sec-10^840_880del^*. Growth and Spk morphology of all the SEC-10 tagged strains and *sec-10* mutants were analyzed. Both the Δ*sec-10* and SEC-10-GFP homokaryon strains had poor colony development, unrecognizable Spk observed by FM4-64 staining LSCM analysis, swollen or bifurcated hyphal tips as revealed by DIC microscopy, and lower growth rate when compared to the wild-type strain. The GFP-SEC-10 protein localized at the hyphal tips much like the C-terminally tagged SEC-10 (Fig. 2) and did not have any hyphal morphology defects or a decrease in growth rate (Fig. 6B; Supplementary video 3). We used transmission electron microscopy (TEM) analysis to examine the cells in greater detail. Representative TEM micrographs of medial sections confirmed the lack of an organized Spk in hyphal tips of the SEC-10-GFP homokaryon strain (Fig. 7). All hyphal tips had large amounts of macrovesicles dispersed in the dome region (Fig. 7 A, B), contrary to the organization seen in previous reports for the wild-type strain, where macrovesicles concentrate primarily in the outer layer of the Spk surrounding a core of microvesicles (Riquelme et al., 2014; Verdin et al., 2009). Interestingly, in few hyphae, a Spk core remained visible (Fig. 7B), whereas other hyphae did not appear to have a defined Spk core as seen in Fig. 7C.1-C.2.

**Figure 7.**
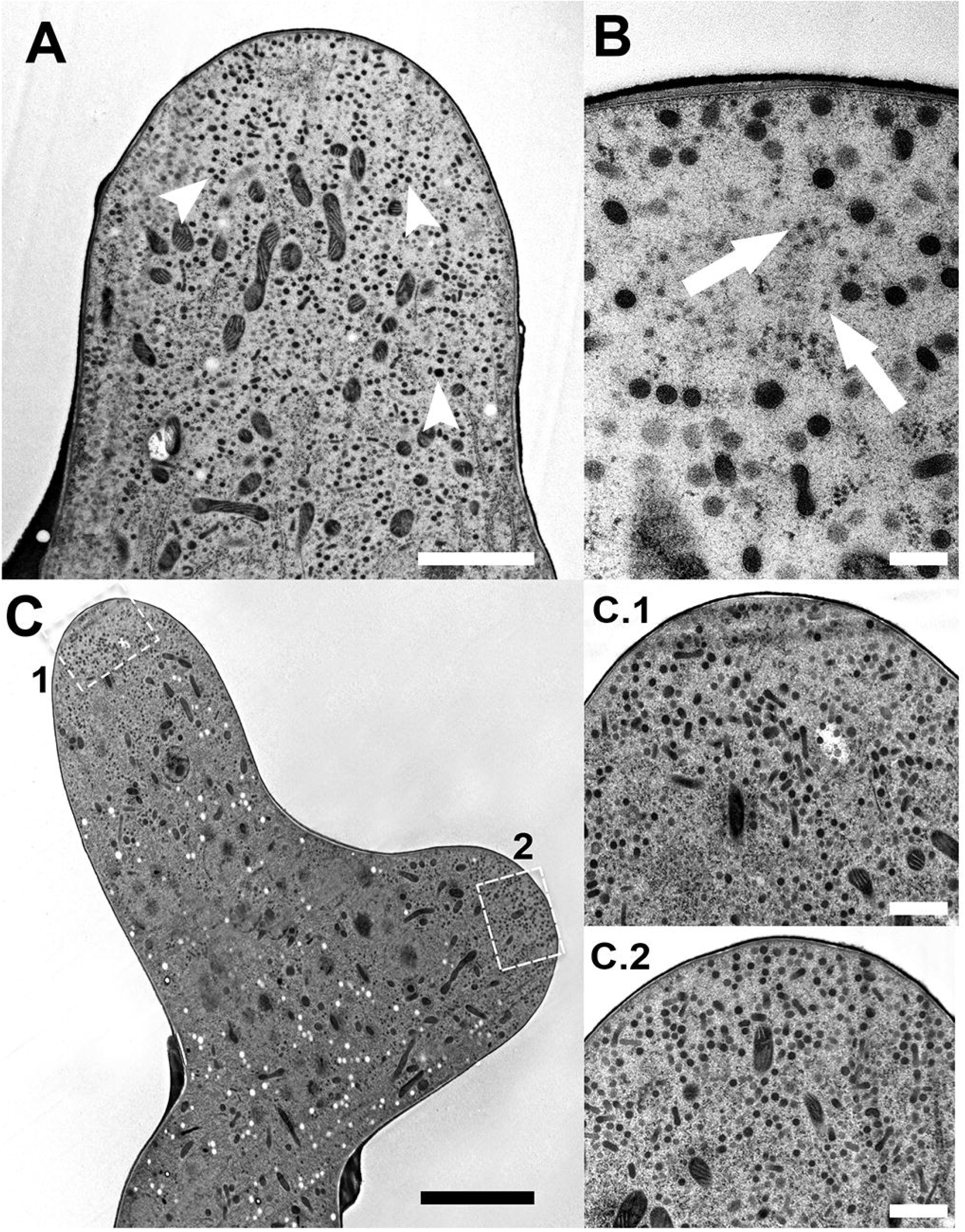
TEM micrographs of hyphal tips of *N. crassa* SEC-10-GFP homokaryon strain (A-B). Tips lacked the characteristic outer layer of macrovesicles and inner layer of microvesicles seen in the Spk. Macrovesicles are seen dispersed in the cytoplasm (arrowheads in A). The SPK core could be clearly seen but lacking an outer core (A) with microvesicles present (arrows in B), while in others a core could not be clearly visualized (C). C.1 and C.2 are magnifications of the dotted line 1 and 2 regions in C. Scale bars A = 2 μm, B = 200 nm, C = 5 μm, C.1 = 1 μm, C.2 = 1 μm.

### SEC-10 co-localizes partially with Spk macrovesicles

The Spk in *N. crassa* has a an inner core of microvesicles that contain chitin synthases (CHS) and an outer layer of macrovesicles that contain glucan synthase (GS) (Sánchez-León et al., 2011; Verdin et al., 2009). In a previous study, the subunits SEC-5, SEC-6, SEC-8, and SEC-15 were shown to localize at the PM of hyphal tips, whereas EX0-70 and EXO-84 localized at the frontal macrovesicular layer of the Spk (Riquelme et al., 2014). SEC-3 was the only subunit that was shown to be present in both locations. To explore whether SEC-10 associates with macro and/or microvesicles, the SEC-10-GFP protein was coexpressed with macro and microvesicle reporters tagged with mCherry fluorescent protein (mChFP). For this, SEC-10 was tagged at the C-terminus with GFP and expressed under the control of *Pccg-1* at the *his-3* locus. As the *sec-10* native locus was unaltered by GFP tagging, the resulting strain did not show any apparent growth defect or affected phenotype. This strain was fused with the strain that expresses CHS-1-mChFP to serve as a reporter for microvesicles, and another strain that expresses GS-1-mChFP as a reporter for macrovesicles. LSCM of the strain expressing SEC-10-GFP and CHS-1-mChFP revealed that SEC-10-GFP did not co-localize with the microvesicles localized at the Spk core (Fig. 8A). In contrast, in the strain expressing GS-1-mChFP and SEC-10-GFP, the SEC-10 protein partially co-localized with macrovesicles on the frontal outer layer of the Spk (Fig. 8B).

**Figure 8.**
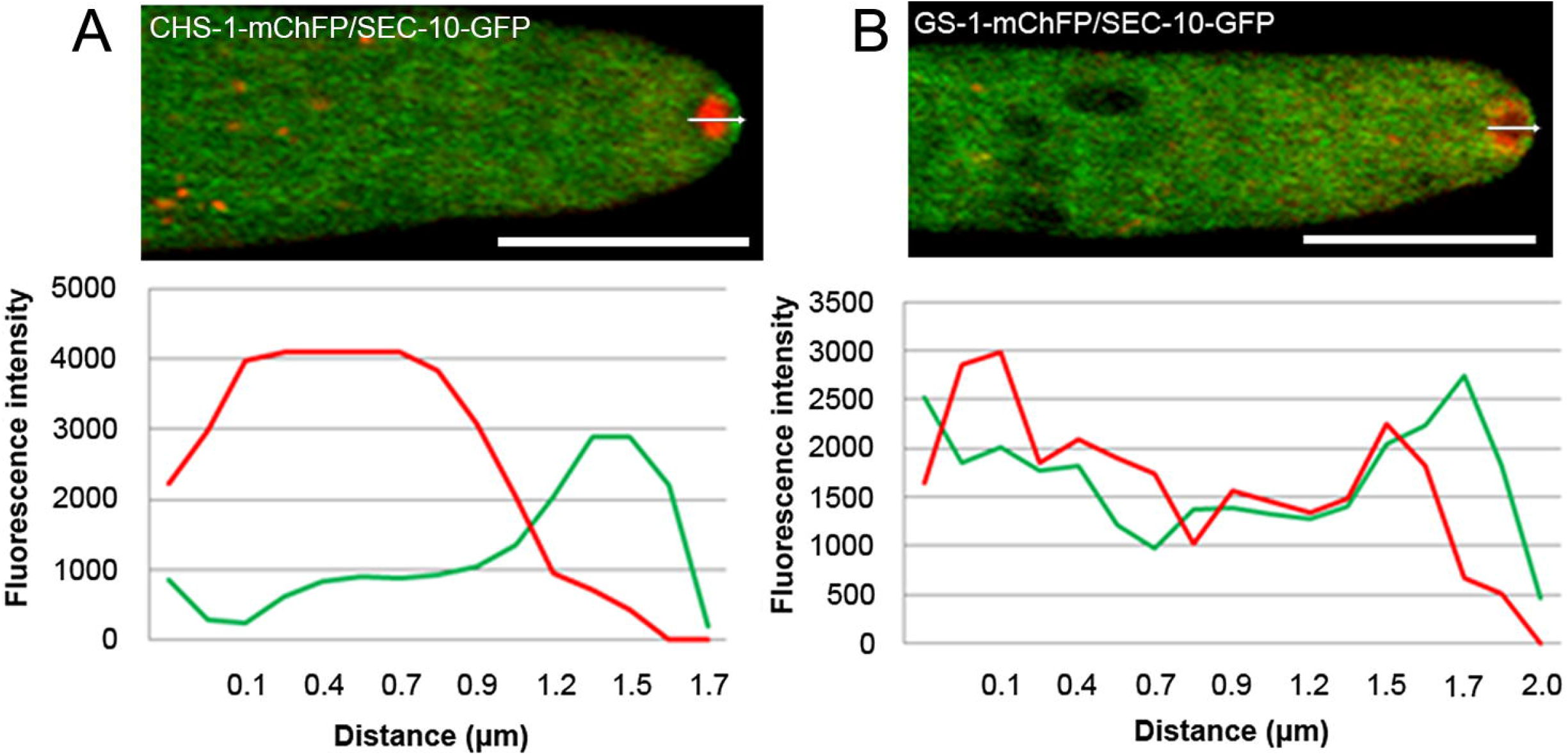
LSCM analysis of strains co-expressing SEC-10-GFP and vesicular markers CHS-1-mChFP or GS-1-mChFP. A) Localization of CHS-1-mChFP in the Spk core. The overlapping signals emitted by GFP and by mChFP reveal that SEC-10-GFP does not co-localize with microvesicles. B) GS-1-mChFP localizes at the outer Spk layer seen as a donut shaped apical body. The overlapping of the signals emitted by GFP and by mChFP reveal partial colocalization. SEC-10 and GS-1 partially colocalize as seen in the fluorescence intensity profile in 2 μm distance from the tip. The white line is the distance analyzed in graph. Scale bars = 10 μm.

### The C-terminus of SEC-10 is important for the interactions with the exocyst subunits

We used pull-down experiments, with anti-GFP nanobody-coupled beads, to examine the assembly and stability of the exocyst complex. SDS-PAGE and Krypton™ staining of the putative interacting proteins purified from the N-terminally and C-terminally tagged SEC-10 strains, indicated that the C-terminal tag severely disrupts the assembly of the exocyst complex. When the C-terminally tagged SEC-10 was pulled down, apparently only one exocyst subunit band was visibly detected in the gel. In contrast, in the N-terminally tagged SEC-10 strain, eight bands could be identified, which corresponded to each subunit of the exocyst complex in *N. crassa,* based on their molecular weight (Fig. 9A).

**Figure 9.**
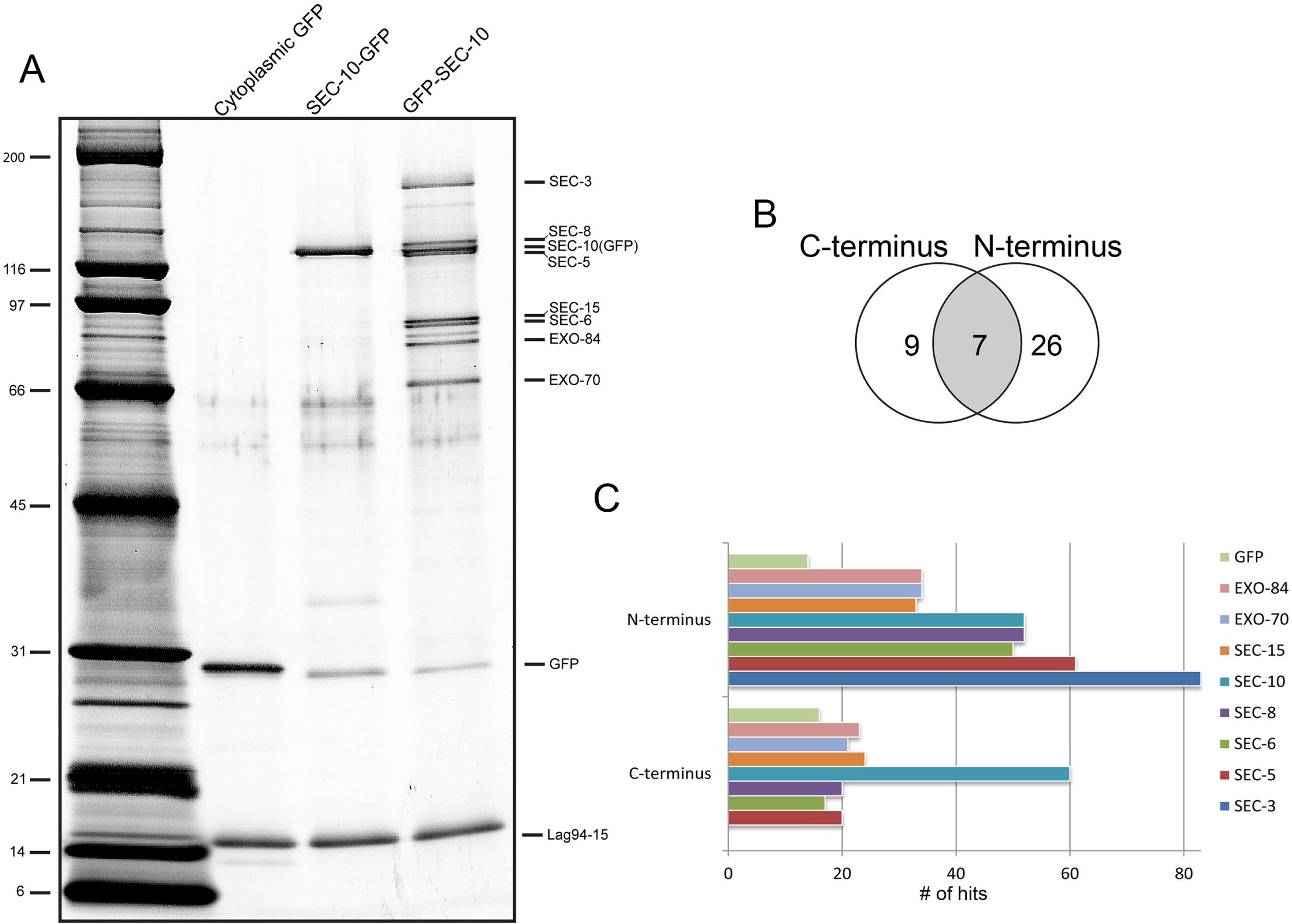
SDS-PAGE and Krypton staining of purified *N. crassa* proteins pulled down by Lag94-15 magnetic beads and unique peptides identified by mass spectrometry. A) The first lane of the stained gel shows cytoplasmic GFP pull-down from a GFP producing strain. Second lane reveals a thick band of SEC-10-GFP and possibly other subunits such as SEC-8 and/or SEC-5. Third lane shows the intact exocyst pulldown by GFP-SEC-10 and Lag94-15. B) Venn diagram depicting the number of proteins detected by mass spectrometry analysis. The left circle of the diagram shows the number of unique proteins identified from the C-terminally tagged sample, the right section corresponds to the N-terminally tagged sample and the intersection contains the proteins shared between both. C) Bar graph comparing the results between the two samples. The X axis shows the number of hits for each of the exocyst subunits while the Y axis shows the unique proteins identified in the N-terminally tagged sample and the C-terminally tagged sample. The subunits are color coded and are placed in the graph in the same order as the key legend.

LC-MS/MS analyses of the pull-downs revealed that the C-terminally tagged SEC-10 had a substantially lower number of bound proteins present when compared to the pull-down from the N-terminal tagged strain. Mass spectrometry analysis of these two samples revealed a total of 258 proteins: 231 proteins in the N-terminally tagged SEC-10 sample, and 107 proteins in the C-terminally tagged SEC-10 sample, with 80 proteins shared by both samples (data not shown). A cutoff of 14 hits or lower was used on the data, and the gene identities were categorized by Gene Ontology (GO) terms (Fig. S3). After the cutoff, only 26 unique proteins remained for the N-terminally tagged sample and 9 proteins for the C-terminally tagged sample, with only 7 shared proteins that corresponded to the exocyst components (Fig. 9B). Regarding the exocyst subunits, the data showed that both samples shared seven exocyst subunits in common, but SEC-3 was excluded by the cutoff from the C-terminally tagged sample. Consistent with the gel in Figure 9A, the C-terminally tagged SEC-10 sample had fewer detected peptides for every exocyst component, suggesting that the GFP tag at the C-terminus leads to weakened affinity of SEC-10 for the rest of the complex. The least abundant subunits, compared to the N-terminally tagged data set, identified in the C-terminally tagged SEC-10 sample were SEC-3, SEC-5, SEC-6, and SEC-8, which represented only around 12-20% of the total amount of each subunit detected in the N-terminally tagged SEC-10 sample. In contrast, SEC-15, EXO-70, and EXO-84 were detected in the C-terminally tagged sample at 28-39% of the subunits detected in the N-terminally tagged sample (Fig. 9C). The number of single peptides for the GFP protein and Lag-94-15 nanobody protein were equally abundant in both samples, which served as internal controls.

## Discussion

Our results demonstrate that tagging of the C-terminal end of SEC-10 disrupted exocyst function and complex stability. This disruption led to aberrant phenotypes observed in SEC-10-GFP homokaryon strains (Fig 6B). Even as a heterokaryon, the SEC-10-GFP strain showed slower growth and wider cells when compared to the wild type. The fact that the SEC-10-GFP heterokaryon has slight growth defects suggests that the tagged SEC-10 proteins can compete with wild type SEC-10 proteins for binding to other exocyst subunits and/or exocyst binding partners.

Strains expressing C-terminally tagged SEC-10-GFP at its native locus under its native promoter, or at the *his-3* locus under the *ccg-1* promoter, revealed localization of the protein at the hyphal tip. However, the endogenously tagged SEC-10-GFP strain had wider and slower growing cells and hyphal deformities including swollen hyphal tips in the homokaryotic state. The ectopically tagged SEC-10-GFP did not show an obvious phenotype, presumably because the native sec-10 locus was intact; this strain still produces a wild type copy of SEC-10 that can assemble into functional exocyst complexes. Furthermore, our LSCM and the TEM micrographs demonstrate that a consequence of exocyst dysfunction is that the hyphae do not contain a conspicuous Spk at their apices. The phenotype observed in this strain is similar to the *sec-5* mutant, which displayed dispersed vesicles and irregular cell shape (Riquelme et al., 2014). Another phenotype of the C-terminally tagged SEC-10-GFP strain was a reduced interseptal distance, which meant that there were an increased number of septa throughout the length of the cell. It is possible that the disorganized Spk and accumulated vesicles at the tip correspond to vesicles that cannot be fused to the PM fast enough to turn over the input of vesicles coming from subapical regions. As more secretory vesicles are being sent to fuse to hyphal tips the vesicles accumulate in the cytoplasm and make the cells wider and swollen at the tips as they wait in queue to be processed by exocyst complexes with apparent inefficiency. This vesicle traffic jam is evidently caused by the unstable exocyst complex that cannot function optimally because the GFP molecule at SEC-10’s C-terminus causes a spatial obstruction with the other exocyst subunits and/or SEC-10 binding partners. The blockage or spatial obstruction caused by the GFP molecule is enough to slow down exocytosis but not enough to block exocytosis entirely. The fact that SEC-10-GFP still localizes at the hyphal tip and that cells are still viable suggest that the exocyst is still working but is lagging in the stage between vesicle docking and vesicle fusion. The slower growth rate of the *Δsec-10* heterokaryon strain suggests that reduced amounts of SEC-10 protein in the cell negatively affected exocytosis and vesicle secretion. This mutant strain could not be obtained in a homokaryotic state, which confirmed that the *sec-10* gene is essential. The low production of conidia in the *Δsec-10* strain could result from a dysfunctional exocyst, since the exocyst has been shown in fungi to play a role in cytokinesis and sporulation (Wang et al., 2016).

What could explain the dysfunction caused by C-terminally tagging SEC-10? Although the C-terminal GFP-tag appeared to disrupt SEC-10 function and incorporation into stable exocyst complexes, a deletion of the C-terminal region in the truncated mutant *sec-10^840_880del^* protein did not have any apparent phenotypic differences. This suggests that the predicted C-terminal disordered region of 40 residues itself does not play an essential role in exocyst function. We initially hypothesized that the C-terminal end of the protein could be important to exocyst function through binding of one or more partner proteins. Upon analysis of the truncated *sec-10^840_330del^* mutant, however, we concluded that SEC-10 binding partners must interact with a larger region of SEC-10, or with residues located more N-terminally. Moreover, the interaction of SEC-10 to assemble into exocyst complexes does not require this C-terminal region; rather, the bulky GFP tag at the C-terminal end of SEC-10 appears to disrupt packing of SEC-10 with other subunits including SEC15 and SEC8, as observed in the S. *cerevisiae* exocyst cryo-EM structure (Mei et al).

Mass spectrometry analyses of the purified C-terminally tagged sample revealed that small amounts of the other seven exocyst subunits could be pulled down by SEC-10-GFP. The Mass spec analysis is sensitive and can detect proteins that the gel could not show. The number of proteins identified was only ~30% of the ones found in the GFP-SEC-10 sample. The amount of SEC-3 that co-precipitated with SEC-10-GFP was so low that it was excluded from the data after the cut-off. This finding suggests that in a C-terminally tagged SEC-10 strain, the exocyst complex is destabilized and SEC-3 does not bind tightly enough to remain bound. However, the other subunits could still bind to SEC-10, albeit with lower affinity or stability. As suggested by the phenotypic analyses, the GFP tag appears to be interfering with protein-protein interactions between SEC-10 and other subunits, leading to a more general disruption and loss of stability of the complex. Deletion of *sec-10* leads to complete disruption of the complex, but in this strain, enough complexes can assemble using the remaining wild-type protein to function in exocytosis, albeit with severe defects in growth and development.

SEC-10 is an essential component of the *N. crassa* exocyst complex, and it localizes at the apical dome in proximity to the plasma membrane. Deleting the *sec-10* gene from the genome is lethal and leads to a mutant phenotype in the presence of a normal copy of *sec-10.* Tagging SEC-10 at the C-terminus disrupts exocyst function when expressed in a homokaryotic state and produces a mutant phenotype. TEM reveals that the C-terminal GFP tagging causes secretory vesicles to accumulate in the cytoplasm instead of in the Spk. Both the heterokaryon *sec-10* knock-out mutant and the SEC-10-GFP homokaryon have growth defects and a Spk that fails to maintain structure during polarized growth at the tip. A fully functional exocyst complex requires all eight subunits to be expressed; with the exception of SEC-5, whose knockout mutant is viable but very affected, deletion mutants for any of the other exocyst subunits are only viable if the strain is a heterokaryon expressing the wild type allele. Expressing SEC-10-GFP in all nuclei (homokaryon) produces a similar cellular phenotype as the heterokaryon *Δsec-10* deletion mutant. The MS data suggests that the C-terminal tag of SEC-10 causes SEC-3 to have decreased affinity for the other exocyst subunits. Although the GFP tag is an artificial construct, its effect on exocyst function is far-reaching, and points to new roles for exocyst that warrant further investigation, including organization of macro- and microvesicles in the Spk, the frequency by which septation occurs along the cell, the continuity of membrane expansion at the tip, and maintenance of cell polarity and growth direction. The future of this work will focus on studying these pathways, as well as the role of SNAREs in exocytosis and explore how they interact with the exocyst complex, which are not fully understood in *N. crassa.* The aim will be to understand the pathways that link the exocyst and the SNARE mediated vesicle fusion.

## Acknowledgments

A. Figueroa-Meléndez was supported by CONACYT research fellowship 638116 and received a PROLAB (Promoting Research Opportunities for Latin American Biochemists program) fellowship through the American Society for Biochemistry and Molecular Biology (ASBMB). We thank the Laboratorio Nacional de Microscopía Avanzada (LNMA-CICESE), Peter Novick from the Department of Cellular and Molecular Medicine, University of California, San Diego, La Jolla, CA and The Mass Spectrometry Facility of the University of Massachusetts Medical School, Shrewsbury, MA, for technical support. We would like to thank Sam O. Obado and Peter C. Fridy from Michael Rout’s lab for technical advice. This work was supported by Consejo Nacional de Ciencia y Tecnología CONACyT, Mexico, grant CB-222375 to M.R., and a grant from the National Institute of Health (GM068803) to M. M.

## Supplementary Videos

Supplementary Video 1. DIC video capture coupled with LSCM of SEC-10-GFP homokaryon strains stained with FM4-64.

Supplementary Video 2. LSCM video capture of the SEC-10-GFP strain showing the characteristic paused growth and moving Spk revealed by FM4-64 staining.

Supplementary Video 3. LSCM video capture of the GFP-SEC-10 strain stained with FM4-64.

**Supplementary Figures**

Supplementary Figure S1. Schematic representation of the methodology used to generate replacement DNA constructs. DNA templates are labeled by gene name and its adjacent UTR whether it is at the 5’ or 3’ end. The PCR products are shown with their respective primers and overhangs. A) C-terminal tagging of SEC-10 using *sec-10*::10×Gly::gfp::hph and hph::3’UTR cassettes. B) N-terminal tagging of SEC-10 using a Pccg::N-gfp fragment that was fused to *hph* and *sec-10* fragments. C) Eliminating *sec-10* by UTR targeting replacement cassettes. D) Eliminating 40 amino acids from the C-terminus of SEC-10 by *hph* selectable gene replacement cassettes that excludes 120 nucleotides in the sequence (red box).

Supplementary Figure S2. One-way ANOVA analysis of hyphal diameters between the different *N. crassa* strains. The strains displayed on the chart from left to right are the 9718 wild type strain, the SEC-10-GFP homokaryon strain (C-terminal tag) and the GFP-SEC-10 homokaryon strain (N-terminal tag). The SEC-10-GFP homokaryon strain had significantly wider cells, compared to the other two strains.

Supplementary Figure S3. Bar graph displaying the total peptides identified in the samples and their Gene Ontology (GO) Categories. A) The X axis shows the number of hits for each of the proteins identified in the samples whereas the Y axis shows the name of the proteins identified in both the GFP-SEC-10 and SEC-10-GFP sample. B-D) The X axis represents the number of different proteins classified under the same GO term and the Y axis shows the name of the GO term. The red colored bars are proteins found in the N-terminally tagged sample and the blue color bars are proteins found in the C-terminally tagged sample.

**Supplementary Tables**

**Supplementary Table 1.** Strains used or generated in this study.

| Strain number | Genotype | Source |
| --- | --- | --- |
| **N1** | *mat a*; WT | FGSC988 |
| **N150** | *mat A*; WT | FGSC9013 |
| **SMRP24** | *mat A*; Δ*mus-51::bar<sup>+</sup>*; *his-3* | FGSC9717 |
| **SMRP25** | *mat a*; Δ*mus-51::bar<sup>+</sup>* | FGSC9718 |
| **SMRP90** | *mat A*; *Pccg-1::chs-1::chfp* | Verdin et al., 2009 |
| **SMRP93** | *mat A*; *Pccg-1::gs-1::chfp* | Verdin et al., 2009 |
| **SMRP410** | *mat A*; *Pccg-1::sec-10::gfp* Heterokaryon | This study |
| **SMRP411** | *mat a*; *Psec-10::sec-10::10xGly::gfp::loxp::hph::loxp*; Δ*mus-51::bar<sup>+</sup>* Heterokaryon | This study |
| **SMRP412** | *mat a*; *Psec-10::sec-10::10xGly::gfp::loxp::hph::loxp*; Δ*mus-51::bar<sup>+</sup>* Homokaryon | This study |
| **SMRP487** | *mat a*; Δ*mus-51::bar<sup>+</sup>*; *sec-10*<sup>2523_2640del</sup>::*hph<sup>+</sup>* Homokaryon | This study |
| **SMRP488** | *mat a*; Δ*sec-10::hph<sup>+</sup>*; Δ*mus-51::bar<sup>+</sup>* Heterokaryon | This study |
| **SMRP489** | *mat a*; *hph<sup>+</sup>::Pccg-1::gfp::sec-10*; Δ*mus-51::bar<sup>+</sup>* Homokaryon | This study |

**Supplementary Table 2.**
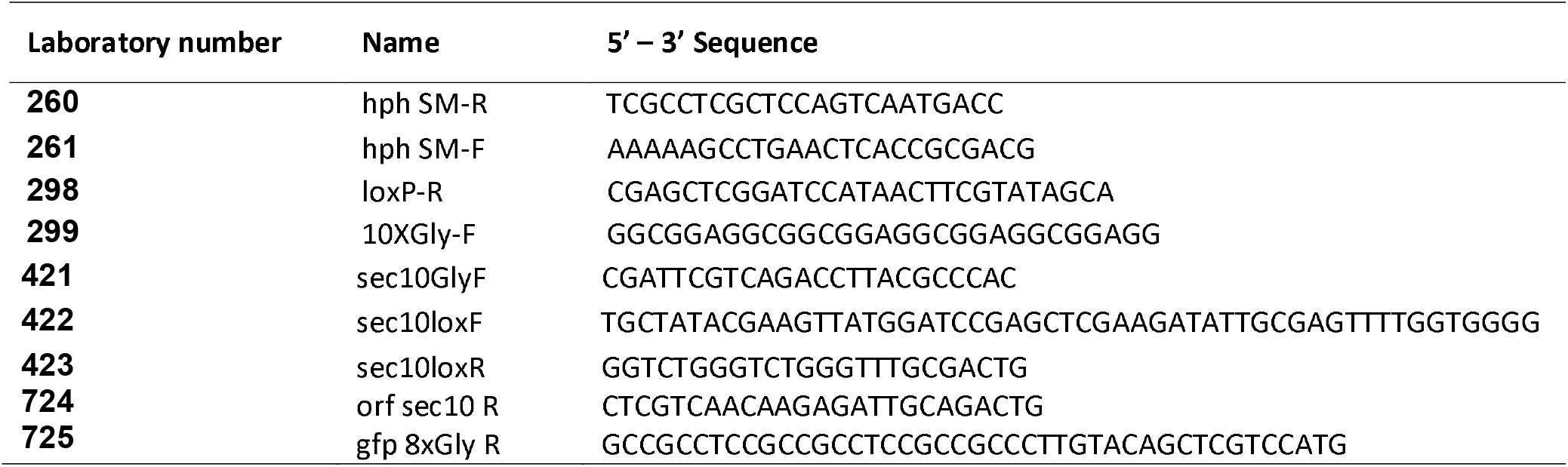
Oligonucleotides used in this study.

