## Supplementary figures and images for "The C-terminal domain of SEC-10 is fundamental for exocyst function, Spitzenkörper organization and cell morphogenesis in *Neurospora crassa*"

### Supplemental Figure 1

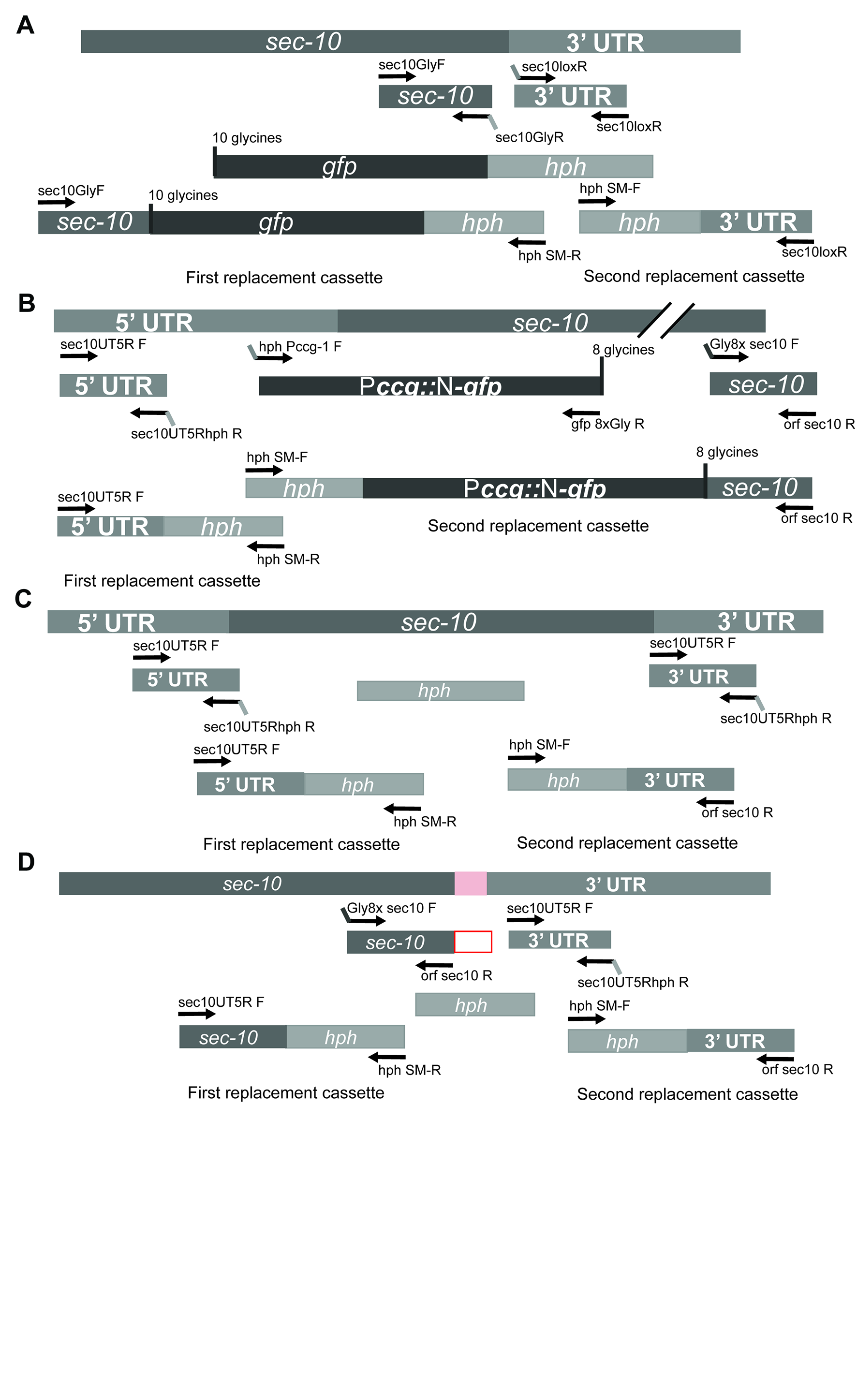

### Supplemental Figure 2

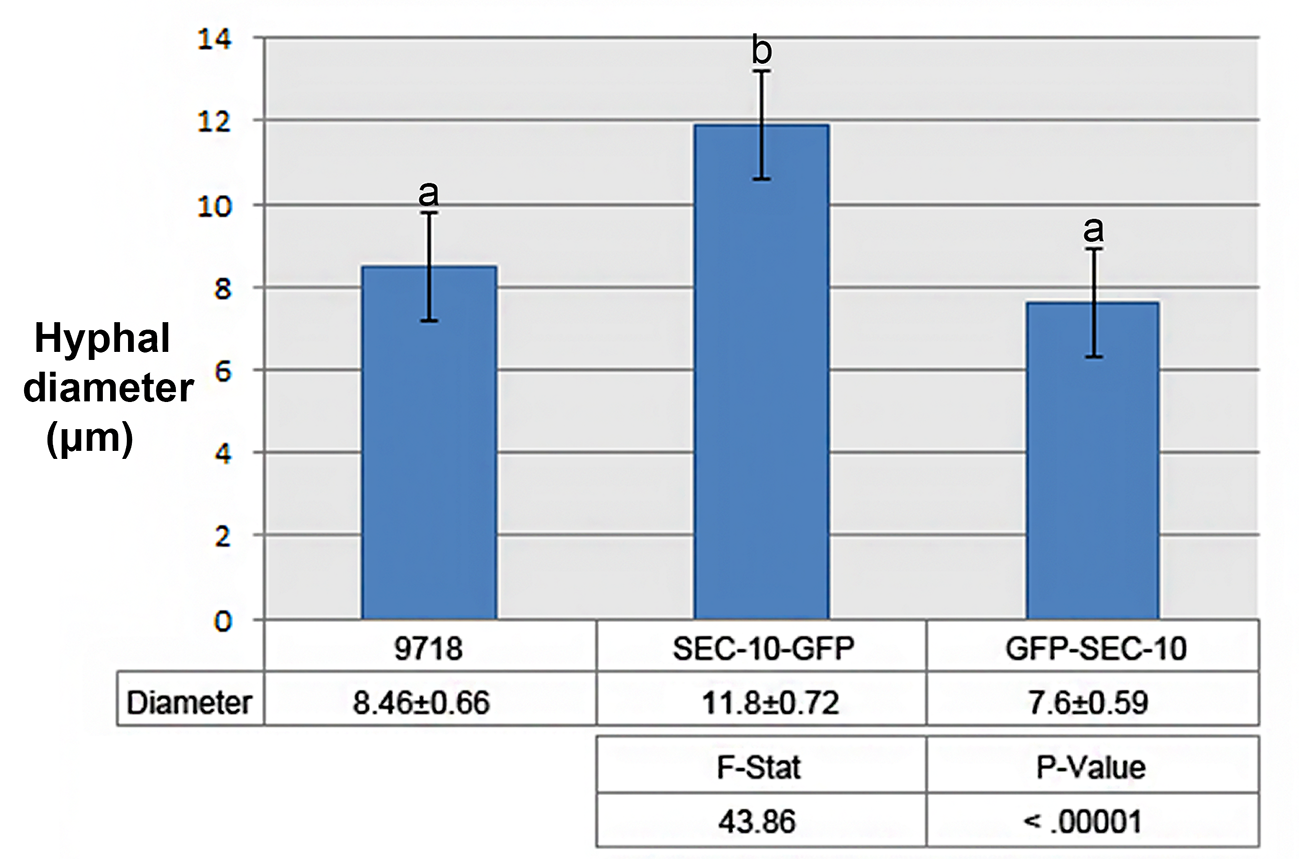

### Supplemental Figure 3

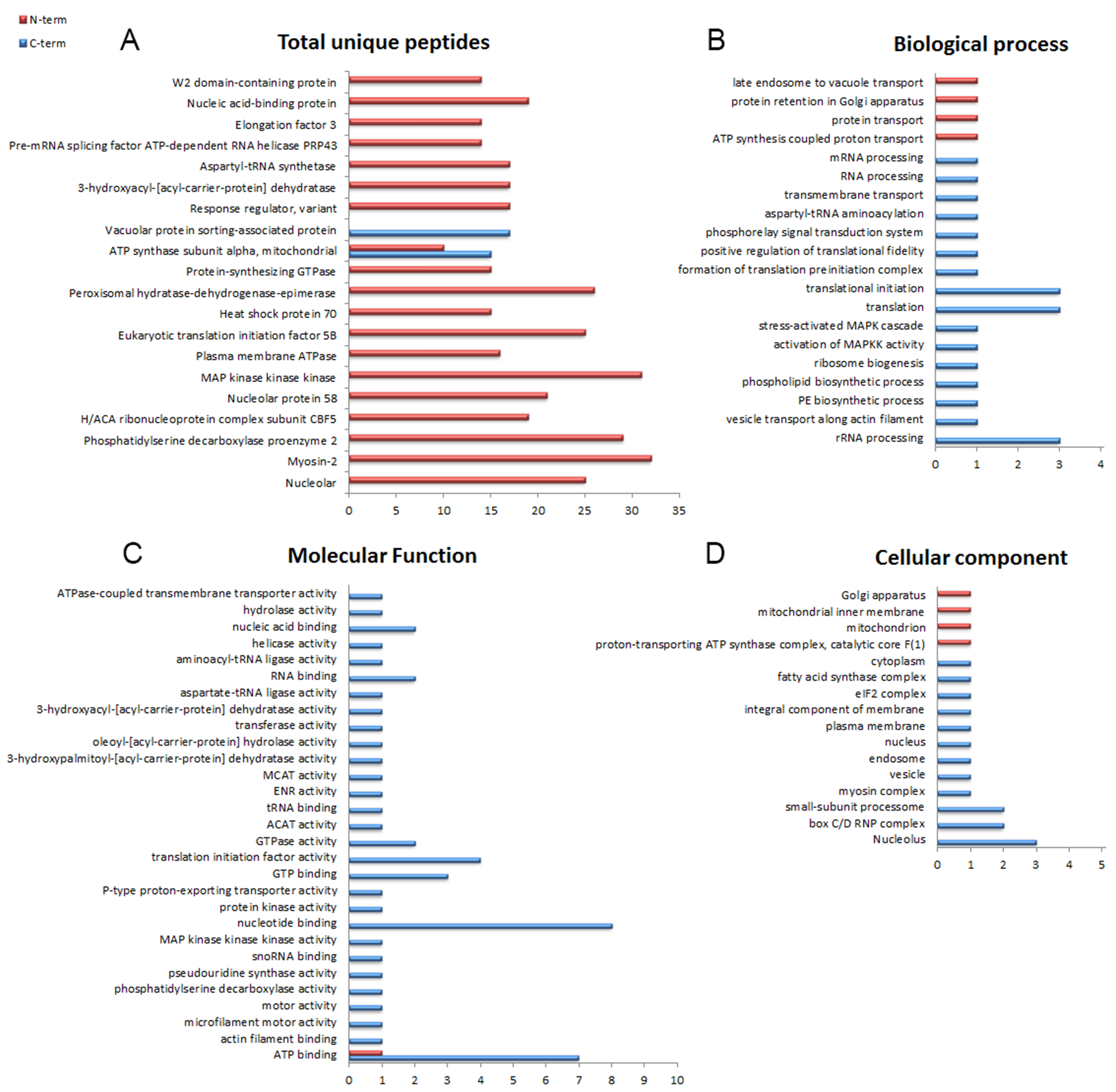
